# iPSC-Derived Ovarian Tissue Restores Ovarian Function in Subfertile Mice and After Gonadotoxic Chemotherapy

**DOI:** 10.1101/697532

**Authors:** K.M. Elias, N.W. Ng, K.U. Dam, A. Milne, E.R. Disler, A. Gockley, N. Holub, G.M. Church, E.S. Ginsburg, R.M. Anchan

**Author notes:** Both authors contributed equally to this publication (K.M. Elias, N.W. Ng). **Raymond M. Anchan, MD, PhD** Division of Reproductive Endocrinology and Infertility, Department of Obstetrics, Gynecology and Reproductive Biology, Brigham and Women’s Hospital, Harvard Medical School, 75 Francis St., Boston, MA 02115.

## Abstract

Many reproductive age women with cancer who receive chemotherapy are exposed to gonadotoxic agents and risk diminished ovarian reserve, sterility, and premature menopause. Previously, we reported the derivation of steroidogenic ovarian cells from induced pluripotent and embryonic stem cells. Derived cells not only produced reproductive hormones, but also displayed markers of ovarian tissue and primordial gametes. Here, we describe that human follicular fluid (HFF), when added to our stem cell differentiation system, enhances the steroidogenic potential of differentiating stem cells and increases the subpopulation of cells that express the ovarian and germ cell markers GJA1 and ZP1, respectively. More importantly, using an *in vivo* model of chemotherapy-induced premature ovarian insufficiency in subfertile nude mice, we demonstrate that orthotopic implantation of these derived cells restores ovarian hormone synthesis and produces functional stem cell-derived oocytes. Additionally, these cells also ameliorate subfertility in nude mice, as demonstrated by the delivery of multiple litters of healthy pups from stem cell-derived oocytes. Collectively, these data support the hypothesis that stem cell-derived steroidogenic ovarian tissue could be used to promote neo-gametogenesis and treat the endocrinologic and reproductive sequelae of premature ovarian insufficiency.

**One Sentence Summary:** We show that orthotopic injection of sorted, differentiated iPSCs in ovaries of subfertile mice restores reproductive hormone synthesis and fertility.

## Introduction

Despite significant and rapid advances in assisted reproductive technologies (ART), few ovarian-sparing options exist for reproductive age women who have cancer and require gonadotoxic chemotherapy *(1)*. Currently, the most reliable method for fertility preservation is pre-chemotherapy fertility treatment cycles using ART to induce the growth of multiple oocytes for cryopreservation, either as oocytes or after fertilization as embryos *(2, 3)*. Unfortunately, referral rates for ART prior to commencing gonadotoxic chemotherapy are low *(4)*. Moreover, compared to similar women without cancer, even women with apparently preserved ovarian function after chemotherapy are more likely to experience infertility as well as reduced success from ART cycles with autologous oocytes *(5, 6)*. This is compounded in this patient population by the fact that mutations in some genes which predispose women to cancer, such as *BRCA1* and Fanconi Anemia pathway members, are also associated with premature ovarian insufficiency *(7, 8)*.

Beyond infertility, premature ovarian insufficiency following chemotherapy treatment causes additional sequelae of estrogen deficiency *(9)*. Cancer survivors experience accelerated bone loss, increased sexual dysfunction, and higher rates of cardiovascular mortality compared to their age-matched peers without cancer *(10–13)*. Restoration of hormones is primarily accomplished by exogenous hormone replacement therapy (HRT); however, there is limited data on the long-term health consequences of prolonged HRT in adolescents and young women, raising concerns about increasing risks of secondary malignancies *(14–16)*. This has led to calls to develop new technologies to preserve or restore ovarian function *(17)*.

In this study, we present a novel approach to the treatment of infertility by generating functional ovarian tissue and oocytes derived from induced pluripotent stem cells (iPSCs). We previously reported that embryonic stem cells (ESCs) and iPSCs may be differentiated into steroidogenic ovarian cells that produce physiologic concentrations of the reproductive hormones estrogen and progesterone *(18–20)*. iPSCs generated from mouse granulosa cells (mouse granulosa cell-derived iPSCs, or mGriPSCs) preferentially differentiate into ovarian cell types due to their epigenetic memory *(18)*. By using patient-specific iPSCs to generate steroidogenic ovarian tissue, these derived cells can be isogenic with the patient. Here, we show that differentiation of mouse iPSCs into steroidogenic and reproductive ovarian cells *in vitro* is enhanced by media containing HFF. Following differentiation, we demonstrate that these ovarian cells and primordial oocytes can be isolated through fluorescence-activated cell sorting (FACS) using a cell surface receptor for a biochemical marker of ovarian reserve, anti-Mullerian hormone receptor 2 (AMHR2). Most importantly, injection of the stem cell-derived AMHR2+ enriched cell population into subfertile mice exposed to gonadotoxic chemotherapy restores ovarian function, as indicated by both the recovery of steroidogenic production and *de novo* generation of stem-cell derived oocytes that display a capacity for fertilization and activation via a calcium ionophore *(21)*. Further, we show that orthotopic injection of differentiated iPSCs into subfertile nude mice restores fertility, as evidenced by healthy offspring derived from these cells. Finally, we propose a central role for the reproductive glycoprotein, Anti-Müllerian Hormone (AMH), in mediating these observations.

## Results

### Differentiated iPSCs regenerate steroidogenic reproductive ovarian tissue *in vitro*

Age-matched normal mouse ovarian tissue (Fig. 1Aa-c) and *in vitro* differentiated mGriPSCs at 2 weeks post-attachment (Fig. 1Ba-c) expressed comparable antigen profiles (Fig. 1Ad-o and Bd-l) for the ovarian and germ cell markers AMHR2, CYP19A1, FOXL2, FSHR, INHB, DAZL, DDX4, ZP1, and ZP2 using immunofluorescent staining. Expression of these markers in the differentiated mGriPSCs was also confirmed by reverse transcription polymerase chain reaction (RT-PCR) (Fig. 1C). Steroidogenic activity in embryoid bodies (EBs) from the differentiated mGriPSCs was confirmed by enzyme linked immunosorbent assay (ELISA), which detected physiological concentrations of estradiol in culture media during extended cell culture (Fig. 1D).

**Figure 1.**
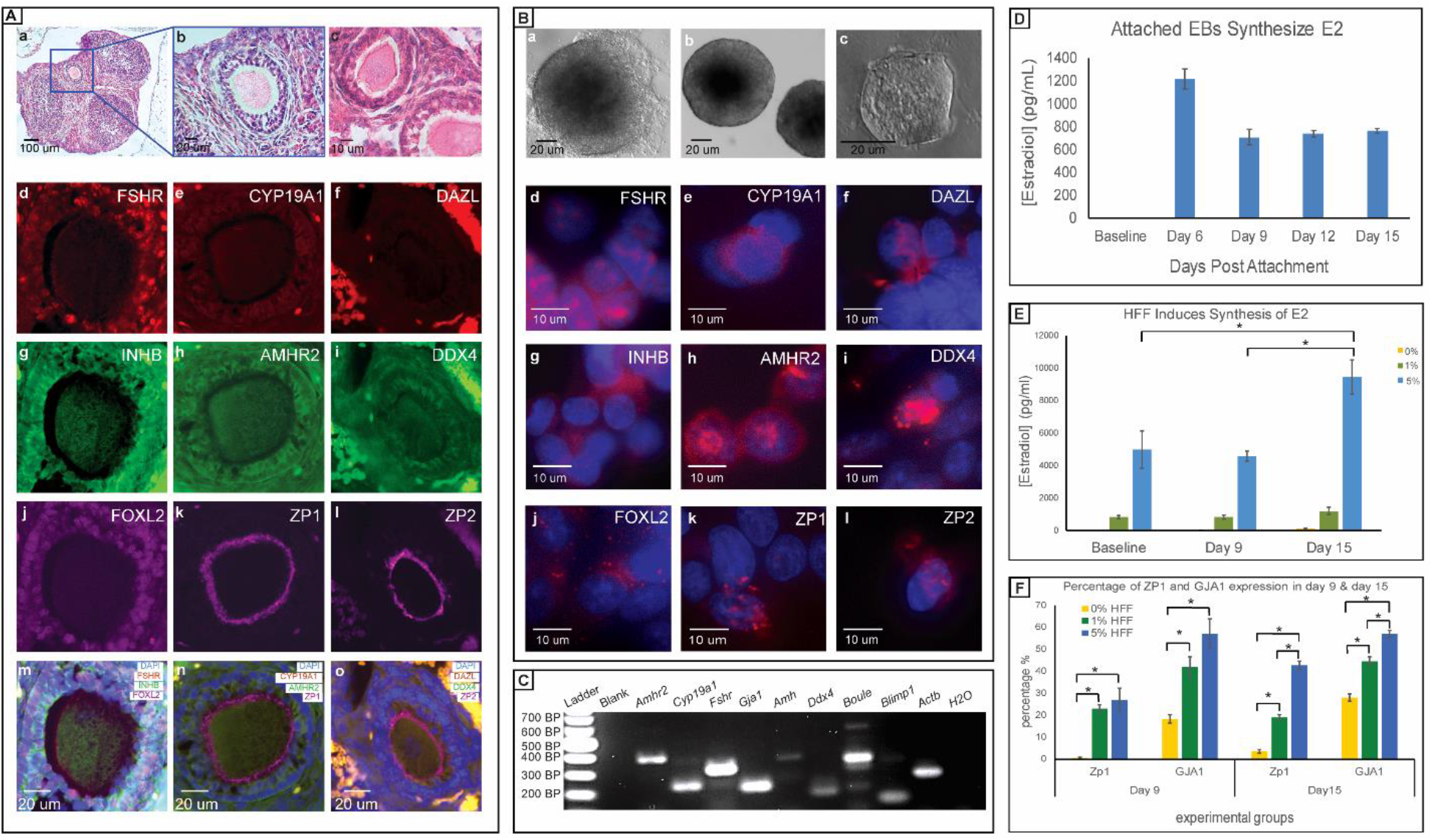
mGriPSCs differentiate into steroidogenic cells that express ovarian and oocyte markers. *In situ* hematoxylin & eosin (H&E) staining of mouse ovarian sections shows follicles (**Aa-c**). Immunohistochemistry (IHC) of mouse follicles demonstrates localization of the ovarian (FSHR, INHB, FOXL2, CYP19A1, AMHR2) and oocyte (ZP1, ZP2, DAZL, DDX4) antigens (**Ad**–**o**). *In vitro* phase-contrast images of a mGriPSC colony (**Ba**), mGriPSC embryoid bodies (EBs) at two weeks post-suspension (**Bb**), and an attached mGriPSC EB (**Bc**). Immunocytochemistry (ICC) shows that differentiated mGriPSC EBs at 15 days post-attachment also express the same corresponding ovarian and oocyte antigens (**Bd-l**). RT-PCR confirms synthesis of these as well as other related ovarian tissue transcripts in these mGriPSCs EBs (*Amhr2*, *Cyp19a1*, *Fshr*, *Gja1*, *Amh*, *Ddx4*, *Boule*, *Blimp1;* **C**). ELISA reveals that differentiated EBs also synthesize the reproductive ovarian hormone, estradiol (E2), at physiologically relevant levels (710-1220 pg/mL) over 15 days (**D**; n=3). The addition of 5% human follicular fluid (HFF) to culture media enhances E2 synthesis in mGriPSCs (**E**; n=3, Mann-Whitney U test, *P*<0.05). 1% and 5% HFF in media also promotes increased expression of an oocyte marker ZP1 (zona pellucida 1) and a granulosa cell marker GJA1^21^(gap junction protein alpha) according to ICC positive cell counts (**F**; n=7, Mann-Whitney U test, p<0.003). Data represented as mean ± SEM.

### HFF promotes differentiation of iPSCs into steroidogenic and reproductive ovarian tissue

mGriPSCs were cultured with media containing 1% or 5% HFF obtained at the time of oocyte retrieval. HFF, collected as pooled samples from multiple patients, was found to contain estradiol (mean > 3,000 pg/mL), FSH (mean = 14.20 IU/L, range = 16.85 IU/L), and LH (mean = 0.26 IU/L, range = 0.27 IU/L). The addition of HFF markedly increased the estradiol synthesis by mGriPSCs over 15 days in culture (Fig. 1E). Additionally, HFF promoted the expansion of a subpopulation of cells expressing the granulosa cell (GC) marker GJA1 *(22)* and the oocyte marker ZP1 (Fig. 1F).

### FACS-purified iPSC-derived ovarian tissue retains endocrine function and ovarian marker expression

After derivation *in vitro*, mGriPSCs were stably transfected with a green fluorescent protein (GFP) reporter using a lentivirus, followed by differentiation into ovarian tissue as previously described *(18, 20)* (Fig. 2A). RT-PCR confirmed that mGriPSCs-GFP retain expression of pluripotency markers (Fig. S1A). These cells were then sorted by FACS using a cell surface level antigen, AMHR2, specific to differentiated ovarian tissue (Fig. 2B) and confirmed by RT-PCR to be expressed in mouse oocytes (Fig. S1B). ELISA revealed preservation of estradiol and progesterone synthesis capacity in both pre- and post-sorted cells (Fig. 2C-E). ICC indicated retained expression of ovarian antigens AMHR2, CYP19A1, and GJA1 in pre-sorted (Fig. 2F-H) and post-sorted cells (Fig. 2I-K). RT-PCR further confirmed these results, as indicated by the retention of RNA transcript expression of ovarian and oocyte-specific genes (*Amhr2*, *Cyp19a1*, *Fshr*, *Gja1*, *Amh*, *Ddx4*, *Boule*, and *Blimp1*; Fig. 2L, M). Sorting mGriPSCs with AMHR2 thus led to a heterogeneous population of differentiated cells that was comprised of both putative GCs and oocytes.

**Figure 2.**
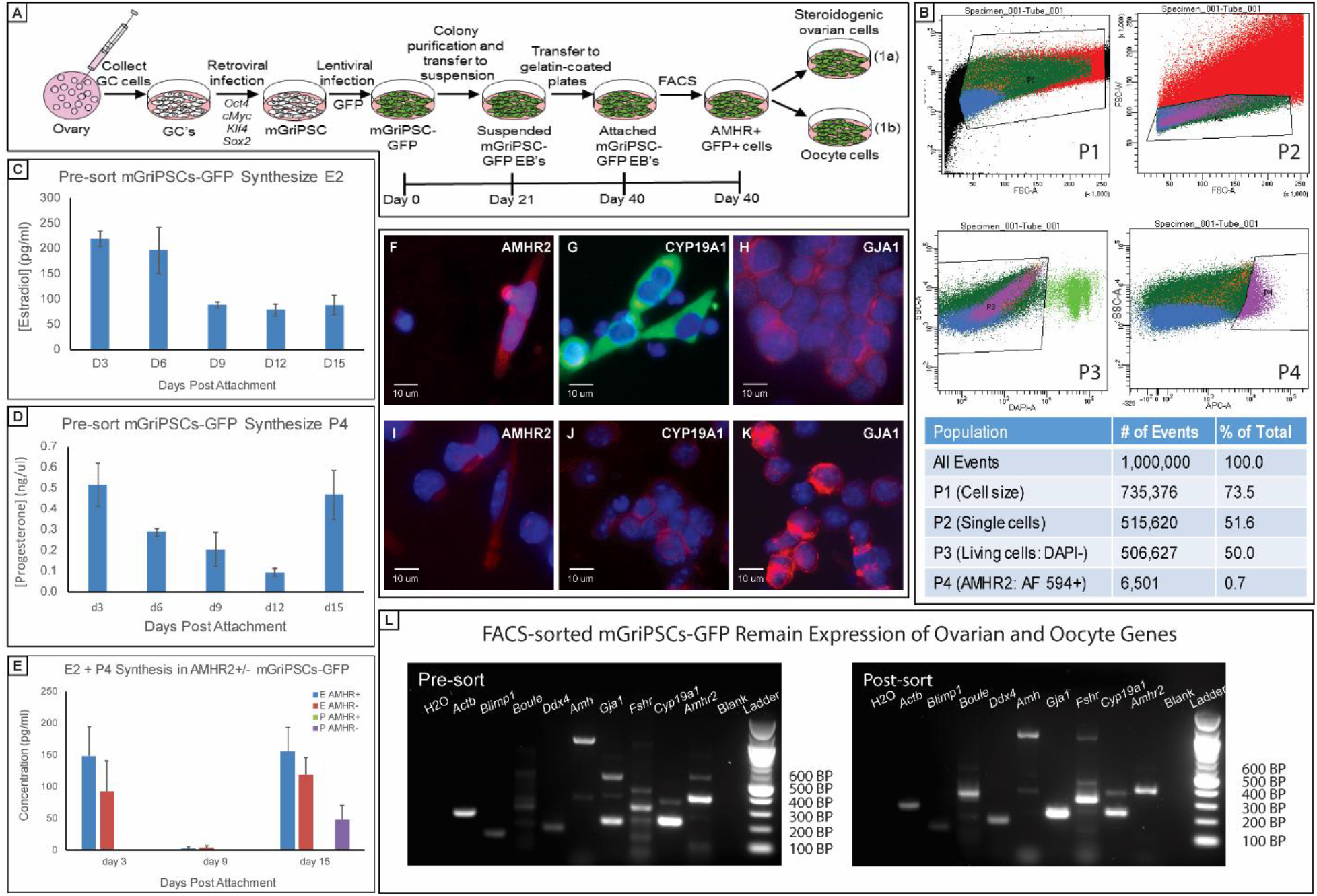
Fluorescence activated cell sorting (FACS) purifies functional iPSC-derived steroidogenic ovarian cells. Schematic representation of the experimental flow for generation of mGriPSCs, green fluorescent protein (GFP) labeling, differentiation, and FACS purification of AMHR2+ mGriPSCs-GFP (**A**). FACS reveals 0.7% of the total population of mGriPSCs-GFP positively expresses ovarian antigen AMHR2 (P4 population; **B**). Pre- and post-sorted mGriPSCs-GFP synthesize E2 and progesterone (P4) through 15 days in culture (**C**, **D**), while post-sorted cells primarily produce E2, consistent with our prior observations *(18)* (**E**). ICC shows that differentiated mGriPSCs-GFP both before (**F**–**H**) and after (**I**–**K**) FACS express the ovarian antigens AMHR2, CYP19A1, and GJA1. The expression of these ovarian markers is further supported by RT-PCR transcriptome analysis of pre- and post-sorted cells (*Amhr2*, *Cyp19a1*, *Fshr*, *Gja1*, *Amh*, *Ddx4*, *Boule*, and *Blimp1*; **L**). For C-E, n=3, data represented as mean ± SEM.

### Alkylating chemotherapy leads to accelerated follicular atresia and breast atrophy

Clinical studies show alkylating chemotherapy results in diminished ovarian reserve with hypoestrogenic effects. In this study, we used athymic nude mice, which are known to experience premature ovarian insufficiency due to age-related rapid follicular atresia *(23)*. To model the additive impacts of exposing a subfertile population to alkylating chemotherapy, nude mice received single intraperitoneal injections of busulfan (12 mg/kg) and cyclophosphamide (120 mg/kg) or 100 ul of vehicle (10% DMSO in PBS; Fig. 3A). Within 63 days of chemotherapy injections, mice exhibited a complete loss of estradiol production (Fig. S2B), in addition to breast atrophy (Fig. S2A). At necropsy, mice that received gonadotoxic chemotherapy displayed smaller ovaries with fewer follicles (Fig. 3Ba) compared to age-matched controls (Fig. 3Bb). This accords with previous mouse studies in which the gonadotoxic effects of cyclophosphamide and busulfan have been demonstrated *(24)*. These mouse studies demonstrated a global loss of ovarian follicles, marginal atrophy, and a presumably increased risk for ovarian failure and infertility after the chemotherapeutic gonadotoxic insult. Therefore, this mouse model resembles the clinical paradigm encountered by reproductive age women undergoing chemotherapy by showing a chemically-mediated decrease in follicle size and number as opposed to a complete elimination.

**Figure 3.**
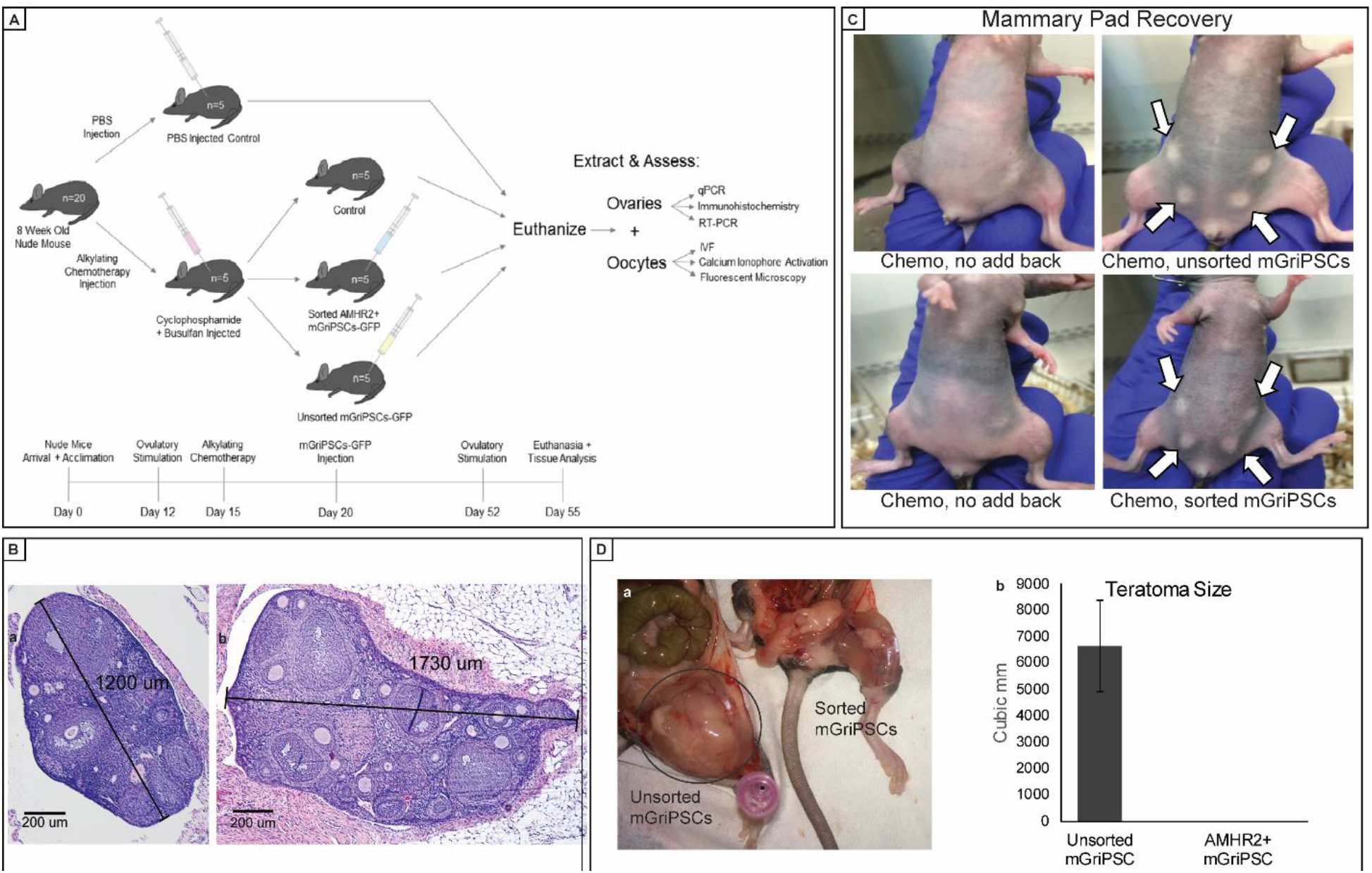
AMHR2+ sorted mGriPSCs-GFP preserve estrogen effects but do not form teratomas. Schematic representation of *in vivo* experimental flow for ovarian injections (**A**). H&E staining revealed partial ovarian atrophy after chemotherapy exposure, as indicated by a decrease in ovary size and fewer follicles (**Ba**) compared to a control ovary (**Bb**). Following alkylating chemotherapy, mice injected intramuscularly with sorted or unsorted mGriPSCs show evidence of mammary pad restoration after 72 hours, whereas mice that did not receive mGriPSCs show no mammary pad recovery (**C**). Injecting FACS-purified cells prevents the formation of teratomas, as compared with unsorted cell injections (**D**; n=3, data represented as mean ± SEM).

### Terminally differentiated AMHR2+ mGriPSCs-GFP restore hormonal function without teratoma formation

We have previously shown that mGriPSCs produce estradiol; however, given the heterogeneity of differentiating EBs, transplantation of these cells may form teratomas *(25)*. We hypothesized that sorting mGriPSCs based on the expression of ovarian cell surface protein AMHR2 might allow selection of a more terminally differentiated cell type, thereby minimizing the risk of tumor formation. Six-week old nude female mice underwent estrous synchronization with pregnant mare serum gonadotropin (PMSG), followed 48 hours later by an ovulation trigger with human chorionic gonadotropin (hCG). Three weeks later, experimental mice received single intraperitoneal injections of busulfan (12 mg/kg) and cyclophosphamide (120 mg/kg), while control mice received 100 uL of vehicle (10% DMSO in PBS). After three additional weeks, experimental mice received intramuscular injections into the left thigh muscle of either vehicle, FACS-purified AMHR2+ mGriPSCs, or unsorted mGriPSCs. Before the stem cell injections, breast atrophy in the chemotherapy exposed mice had been noticeable (Fig. 3C), indicative of estrogen deficiency. Within 72 hours of stem cell injection, the breast atrophy had been reversed in the mice receiving sorted or unsorted cells (Fig. 3C), suggesting restoration of estradiol production. Mice were followed for an additional one month. While all mice that received unsorted cells developed large teratomas, no macroscopic or microscopic teratomas were seen in mice that received sorted cells (Fig. 3D).

### iPSC-based *de novo* generation of steroidogenic GCs and mature oocytes *in vivo*

While intramuscular injections of AMHR2+ mGriPSCs appeared to preserve hormonal function in chemotherapy treated mice, because of the absence of tumor formation, we were not technically able to retrieve the cells for subsequent analysis. Therefore, we next investigated the effect of orthotopic injection of mGriPSCs-GFP. As in the prior experiment, mice underwent estrous synchronization and then received alkylating chemotherapy. Rather than allowing the follicles to atrophy completely as in the prior experiments, we planned a shorter time interval for stem cell injections to model a rescue therapy. Therefore, the following week, mice received direct intraovarian injections of GFP-labelled unsorted or sorted AMHR2+ mGriPSCs via laparotomy into the left ovary. Shortening the timeframe between chemotherapy and stem cell administration also made it easier to perform surgical manipulation of the ovaries before they became too atrophic. Mice were then followed for one month. Mice then underwent a second 48-hour cycle of PMSG/hCG ovarian hyperstimulation, followed by euthanasia.

Oocytes were then collected directly from the ovaries by puncturing the bursa *ex vivo* and flushing the oviducts. Mature oocytes collected from mice injected with sorted mGriPSCs-GFP not only expressed GFP under fluorescent microscopy (Fig. 4A,B; Fig. S3A-M), but also displayed evidence of both calcium ionophore activation (Fig. 4C; Fig. S3J-M) and fertilization (Fig. 4D; Fig. S3D-I). The *in-situ* (Fig. 4E) and *ex vivo* (Fig. 4F) expression of GFP in GCs of mGriPSCs-GFP treated mice suggests *de novo* generation of stem cell-derived ovarian tissue, as well as the oocytes themselves. Ovarian follicle immunosections from mice injected with sorted cells also expressed GFP, which co-localized with AMHR2 and DAZL (Fig. S3N,O). Additionally, quantitative reverse transcription PCR (RT-qPCR) indicated increased *in vivo* expression of terminal oocyte marker *Zp1* in the ovaries that received the sorted mGriPSCs-GFP injection compared to the contralateral uninjected ovaries or the ovaries of mice that received chemotherapy but no stem cell injections (Fig. S4). After gonadotropin hyperstimulation, 18 mature oocytes were recovered from the mouse ovaries that were previously orthotopically injected with sorted cells, whereas five oocytes were collected from these mice’s contralateral ovaries in which no cells were injected. Of these 18 retrieved oocytes, eight expressed GFP (Fig. 4G; Fig. S3), indicative of their stem cell origin. This suggests neogametogenesis, as well as salvation of native oocytes through paracrine effects. RT-PCR analysis on these oocytes showed co-expression of *Sry* and *Gfp*, confirming both successful fertilization and derivation from the mGriPSCs-GFP lineage (Fig. 4H). No oocytes were collected from the left ovaries of the mice injected with unsorted cells, as large teratomas obstructed the collection process (Fig. S5).

**Figure 4.**
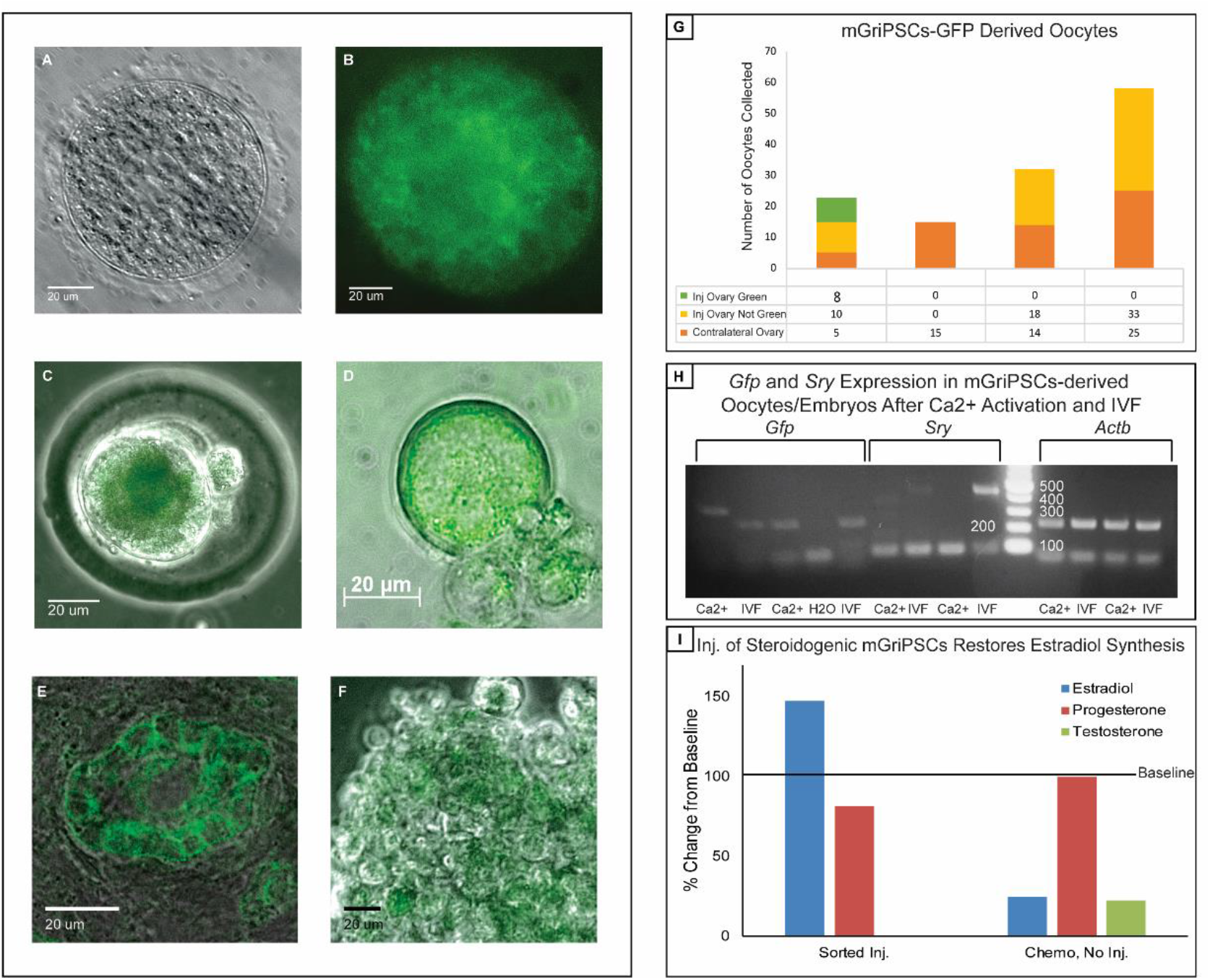
Neogametogenesis with differentiated iPSCs *in vivo*. Images A-F depict cells and tissues collected from mouse ovaries exposed to gonadotoxic chemotherapy followed by injection of sorted mGriPSCs-GFP. Corresponding phase contrast (**A**) and fluorescent microscopy (**B**) images of a mature oocyte expressing GFP. Maturation and functionality of GFP-labeled oocytes is supported by the ability to activate with calcium ionophore (A23817; **C**) and fertilize *in vitro* (**D**). IHC indicates localization of GFP to granulosa cells of the follicle (**E**). Isolated granulosa cells also express GFP (**F**). Oocytes with and without GFP expression are counted from mice in all treatment groups, in both the ovary injected with stem cells and the contralateral ovary (**G**). RT-PCR transcriptome analysis confirms *Gfp* expression in calcium ionophore activated oocytes and IVF embryos, as well as *Sry* expression in IVF embryos (**H**). Compared to mice that receive chemotherapy without mGriPSCs injections, mice that receive mGriPSCs injections after chemotherapy show an increase in blood E2 levels (**I**; n=3, data represented as mean percent change in hormone concentrations before and after mGriPSCs injections, as measured by ELISA). As previously reported, since E2 is the predominant reproductive hormone synthesized by differentiating mGriPSCs, P4 is observed to remain unchanged under all conditions^17^. Testosterone is only detectable in mice that receive chemotherapy but no mGriPSCs injections (**I**).

Injection of differentiated AMHR2+ mGriPSCs-GFP into the ovary yielded an increase in estradiol production from pre-to post-stem cell injection conditions (Fig. 4I). Further, in alignment with our prior studies, the progesterone synthesis between different experimental groups was not significantly different (Fig. 4I), as mGriPSCs appear to preferentially differentiate into estradiol synthesizing GCs, presumably the consequence of their epigenetic memory. ELISA detected testosterone production in only the group that received alkylating agents without stem cell injections (Fig. 4I). Testosterone measurements support the hypothesis that observed differences between groups in estradiol levels, which is synthesized from testosterone via aromatase, are not restricted by availability of substrate. Together, this shows that AMHR2+ mGriPSCs-GFP can produce both functional GCs and mature oocytes.

### Orthotopic injection of AMHR2+ mGriPSCs-GFP enhances fertility in nude mice

As lymphodepletion improves the success of bone marrow stem cell engraftment, we considered the possibility that pretreatment with chemotherapy might similarly deplete native follicles to allow engraftment of the mGriPSCs or that the chemotherapy itself might otherwise somehow modify the ovarian microenvironment. To exclude this possibility, we injected stem cells into the ovaries of subfertile mice not exposed to chemotherapy. Five weeks after injecting sorted AMHR2+ mGriPSCs-GFP into the ovaries of five nude mice, two of the nude mice were hyperstimulated with PMSG and hCG and euthanized for oocyte retrieval and *in vitro* fertilization (IVF). Six oocytes were retrieved from mouse number 1 and seven oocytes were retrieved from mouse number 2. Eight out of 13 of these oocytes expressed GFP according to fluorescent microscopy. All of these oocytes exhibited the capacity to fertilize and develop into at least two-cell stage embryos when exposed to mouse sperm (Fig. S3).

Eight weeks after injecting the sorted AMHR2+ mGriPSCs-GFP, the remaining three female nude mice were mated with wild type Black 6 males. All three mice became pregnant within two weeks of housing with males (Fig. 5A). Two of the pregnant mice were euthanized around embryonic day 18. Mouse number 1 produced three embryos and mouse number 2 produced six embryos, all of which were morphologically normal (Fig. 5B). Confocal microscopy used on freshly harvested embryos directly after euthanasia showed that various translucent embryonic tissues expressed GFP (Fig. 5C), in addition to the chorionic villi at the placental implantation site on the dam’s uterine cavity (Fig. 5D). Mouse number 3 became pregnant, displayed normal gestation, and delivered a litter of seven pups, three of which expressed GFP according to fluorescent IHC (Figure 5A, J-M). Mouse number 3 then became pregnant two additional times and delivered litters of nine and five pups, respectively, within the span of two months. All pups (F1 generation) were healthy at birth and displayed normal development (Fig. 5E). According to fluorescent IHC (Fig. 5F-I), five out of nine of the embryos from the first two litters (mouse 1 and 2) expressed GFP, indicating that they were derived from the injected mGriPSCs-GFP. IHC also verified that seven of the sixteen pups from mouse number 3’s first two litters expressed GFP (Fig 5J-M). Four F1 female pups were mated with two F1 male pups. All four females gave birth to litters between 9 and 11 pups. This total of 41 pups represents the F2 generation of mice from the original injected mGriPSCs (Fig. S6).

**Figure 5.**
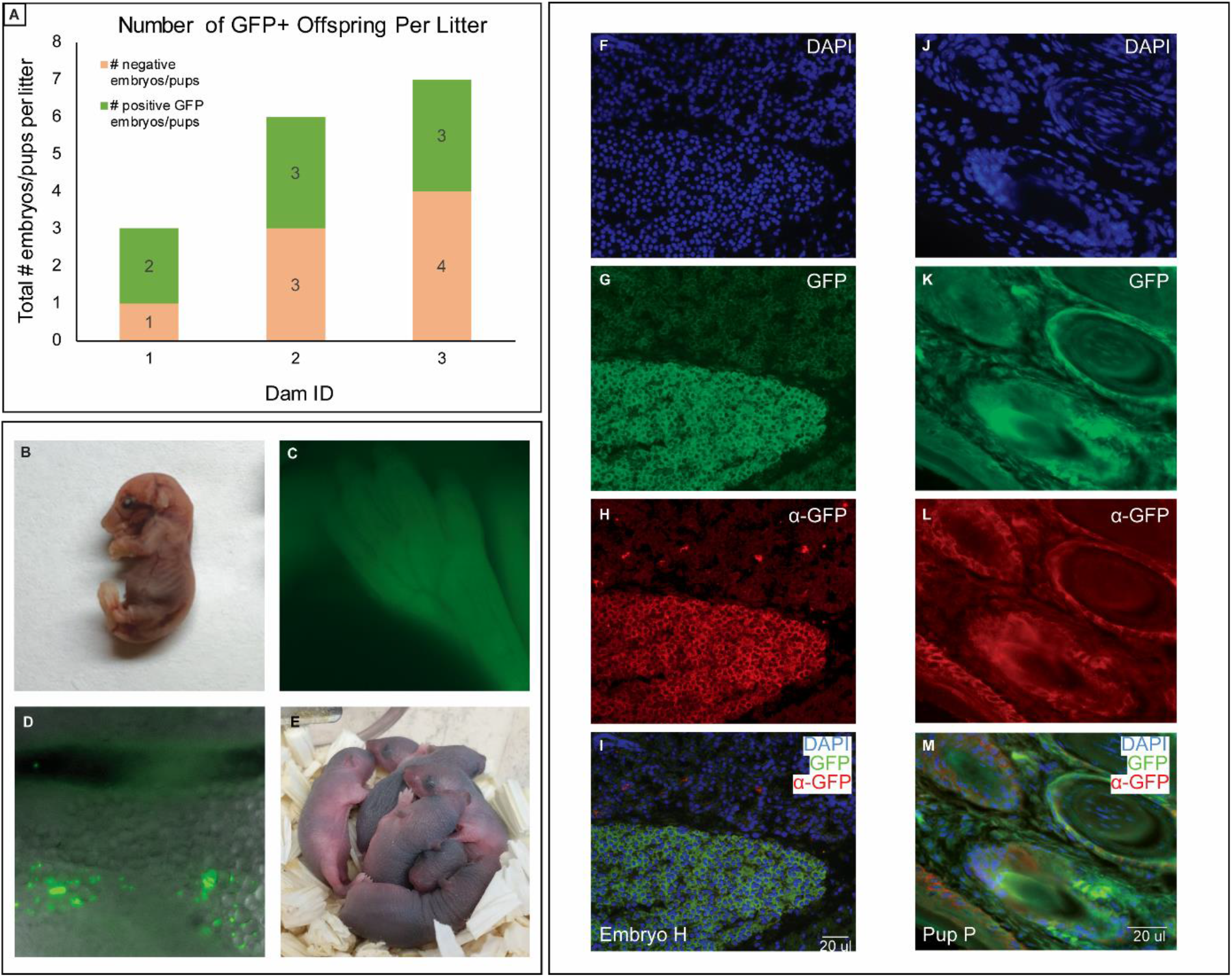
AMHR2+ mGriPSCs-GFP restores fertility in nude mice. Breeding of all three nude mice results in GFP-labeled, chimeric offspring (**A**). Embryo gross morphology at embryonic day 18 (**B**) appears normal. Confocal microscopy demonstrates endogenous expression of GFP in the paw (**C**) of an E18 embryo, as well as at the placental implantation site where the chorionic villi adhere to the mother’s uterine cavity (**D**). Pups were carried for full term and displayed normal development (**E**). Fluorescent IHC on whole embryo **(F-I)**and pup tail snip **(J-M)**sections reveal expression of endogenous GFP (**G, K**), as well as corresponding immunostained GFP (**H, L**) and merged images (**I**, **M**), confirming the derivation of these chimeric embryos from GFP-labeled stem cells.

### AMH synthesis is observed in stem cell treated mice

AMH is typically observed in the native ovary and is a marker for ovarian follicular reserve *(27)*. The differentiated mGriPSCs-GFP that were sorted and later injected into mouse ovaries demonstrated AMH synthesis *in vitro* (Fig. S7 as well as in the tissue (Fig. 6A,B) and serum (Fig. 6C) from mice treated with mGriPSCs-GFP. As expected, chemotherapy accelerated the reduction in AMH levels compared to control mice (Fig. 6C). In contrast, AMHR2+ mGriPSCs preserved AMH levels *in vivo*, an effect not seen with unsorted mGriPSCs (Fig. 6C). Interestingly, when we compared *Amh* mRNA levels between the left ovary, which received AMHR2+ mGriPSCs-GFP, and the right ovary, which did not, *Amh* expression was high in both ovaries compared to chemotherapy exposed ovaries from mice that did not receive mGriPSCs (Fig. 6D). This could reflect paracrine effects mediated by the injected stem cells.

**Figure 6.**
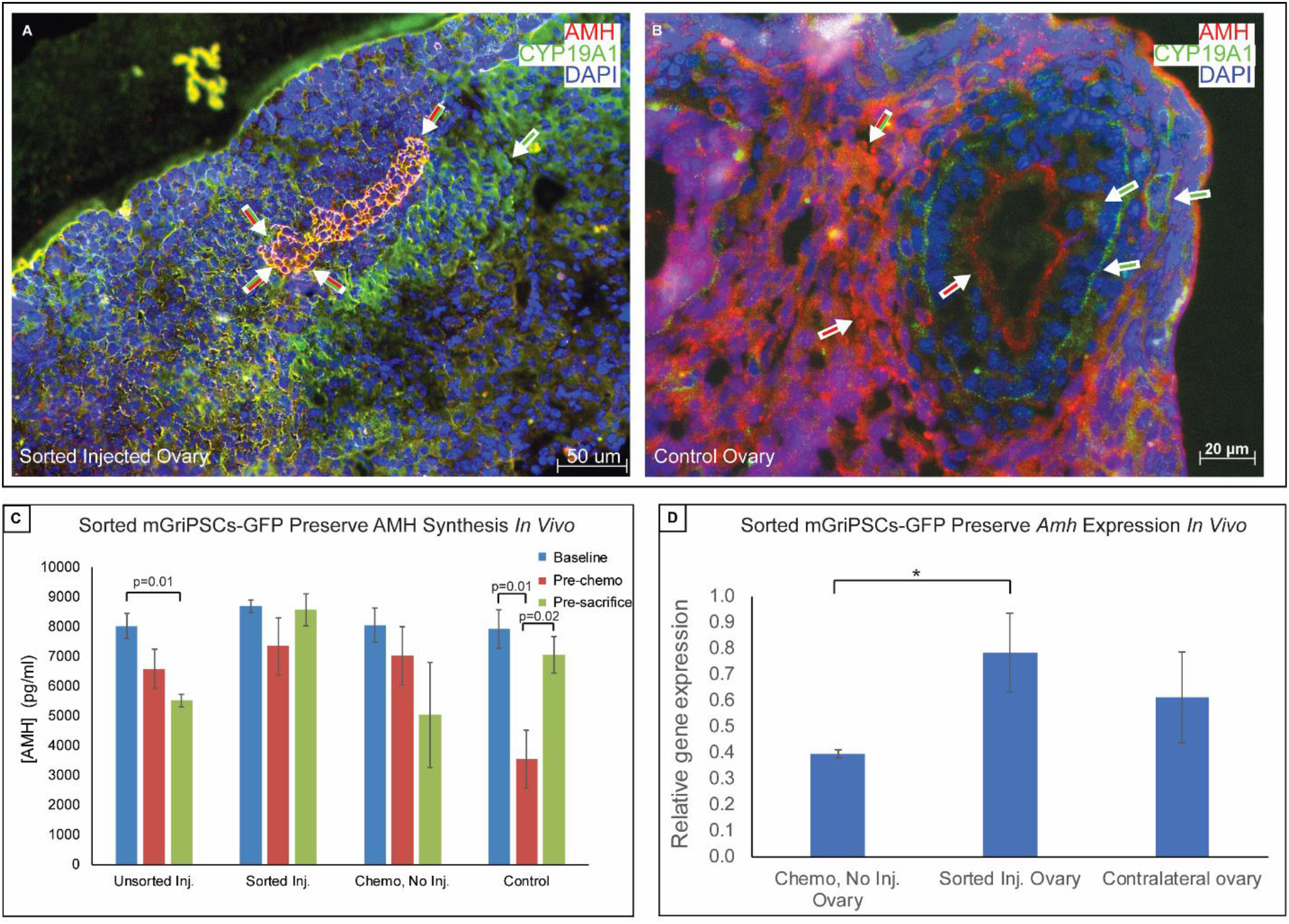
Sorted mGriPSCs-GFP synthesize AMH *in vivo*. IHC of an ovary exposed to gonadotoxic chemotherapy and injected with sorted mGriPSCs-GFP confirms estradiol and AMH synthesis (**A**), as compared with a control ovary (**B**). Mice exposed to chemotherapy show reduced AMH synthesis, but those that receive sorted mGriPSCs-GFP injections retain AMH synthesis after stem cell injections (**C**; n=3, Mann-Whitney U; data represented as mean ± SEM, as measured by ELISA). RT-qPCR analysis on RNA extracted from ovaries exposed to chemotherapy reveals greater expression of *Amh* in mice injected with sorted cells compared with those that were not injected with stem cells (**D**; n=3, Mann-Whitney U, p<0.05; data represented as mean gene expression fold change ± SEM, as compared with control ovary gene expression).

## Discussion

In this study, we report a potential autologous iPSC system for the *de novo* generation of functional oocytes and fertility restoration. While early studies have shown differentiation of murine embryonic stem cells to express primordial germ cell markers *(28)*, the capacity to mature these cells *in vitro* and to fertilize them has been difficult. Recent observations suggest that the ovarian cortex plays a central role in mediating the maturation and development of primordial germ cells *(29, 30)*. The ovarian cortex is comprised of stroma and functional follicles. The primary follicular cell types include mural and cumulus GCs that line the antrum of the follicle, which also contains the primordial oocyte. The primary endocrine cell types of the ovary are the thecal cells, which make androgens, and the GCs, which produce estrogen and progesterone from androgen precursors.

The current study builds upon our prior report demonstrating the homotypic differentiation of mGriPSCs into steroidogenic ovarian tissue *(18, 20)*. While the feasibility of generating stem cell-derived oocytes using reconstituted ovarian tissue has been described *(18)*, our work for the first time shows that oocyte development and the supporting ovarian cortical matrix can both be promoted directly from induced pluripotent stem cells *(31, 32)*. A key observation in our studies is the capacity for dfferentiated, sorted mGriPSCs to contribute to the *de novo* generation of oocytes, as well as to form ovarian follicles as evidenced by the GFP-labelled follicular granulosa cells. What remains unclear after FACS purification is whether: 1) two distinct types of AMHR2+ precursors arise that lead to either oocyte or GC lineages, or 2) a common, AMHR2+ stem cell-derived precursor arises that later differentiates into either oocytes or GCs. Of note, RT-PCR analysis of native mouse oocytes revealed co-expression of AMHR and ZP1 (Fig. S1B); therefore, the generation of two cell types arising from AMHR+ sorted cells seems highly plausible and supports the notion that mGriPSCs are regenerating ovarian tissue.

*In vivo*, the developing primordial oocyte is bathed in follicular fluid. This close association of HFF with the oocyte supports an important function for HFF in oocyte maturation *(33–35)*. We found that differentiation into oocytes from mGriPSCs was enhanced by using HFF, analogous to what one observes *in vivo.* The resulting differentiated stem cells synthesized autologous steroid hormones and expressed normal markers of germ cells and granulosa cells.

In humans, gonadotoxicity of chemotherapeutic agents is highly variable and dependent on dose, patient age, and prior ovarian reserve. Dosing related toxicity is influenced by absolute and cumulative dosing. Nearly 20% of breast cancer cases present in women younger than 50 years old *(36)*. While newer treatment regimens employed are less gonadotoxic, regimens still consist of combination medications that include cyclophosphamide, known to deplete the number of primordial follicles, thereby potentially leading to infertility *(37)*.

Typically, during a menstrual cycle, the hypothalamic-pituitary-ovarian axis coordinates the release of gonadotropins to recruit ovarian follicles, initiating the concomitant development of oocytes *(38)*. As such, during controlled ovarian hyperstimulation, injections of gonadotropins (FSH/LH) recruit ovarian follicles and increase estradiol synthesis *(39)*. We observed that gonadotropin-induced ovarian hyperstimulation resulted in an appropriate physiologic response with maturation of follicles and gametes further supporting the conclusion that the stem cell-derived ovarian cells exhibit normal endocrine behavior comparable to native ovarian cortex. Furthermore, we have shown that the orthotopic injection of differentiated, AMHR2+ iPSCs into nude mice both with or without chemotherapy facilitated the *de novo* generation of stem cell-derived oocytes. This is demonstrated by the presence of GFP+ oocytes in both groups of nude mice. Importantly, the ability to fertilize and use a calcium ionophore to activate these stem cell-derived oocytes demonstrates functional neo-gametogenesis. This functionality is further supported by delivery of multiple litters of iPSC-derived. Homozygous nude mice are known to be subfertile, as they exhibit ovarian follicular atresia, low oocyte counts, and significantly diminished synthesis of gonadotropins and reproductive hormones such as estradiol and progesterone *(40–42)*. In a previous study, fewer than 10% of nude mice were able to achieve pregnancy and deliver pups when mated with wild type males *(40)*. In our experiments, we demonstrated that orthotopic injection of differentiated AMHR2+ mGriPSCs-GFP and subsequent breeding with wild type males led these subfertile mice to not only become pregnant with sizeable litters, but also to repeatedly deliver healthy pups that express GFP.

Interestingly, injection of these AMHR2+ mGriPSCs-GFP into a single ovary of nude mice following chemotherapy also improved endocrine function in the contralateral ovary. The capacity to rescue the contralateral ovary suggests this is mediated by a paracrine mechanism. This agrees with observations from other groups that describe the potential role for paracrine mechanisms mediated by injecting bone marrow and mesenchymal stem cells for chemoprotection of ovaries *(43–45)*. The precise paracrine mechanism of action remains elusive for those stem cells; however, AMH is a promising candidate molecule given its chemoprotective and preventative effects on follicular atresia and apoptosis *(36–38)*. AMH is exclusively synthesized by granulosa cells during the early stages of follicular development *(49)*, and AMH serum concentration is an important measure of ovarian reserve *(27)*. Additionally, the effect of gonadotoxic agents on ovarian follicular development is reflected by lower serum concentrations of AMH *(50, 51)*. Moreover, low AMH levels are prognostic for poor probability of spontaneous conception as well as impaired response to ART *(52)*. Post-chemotherapy AMH levels are also an important biomarker for early onset of menopause *(53)*. As such, we explored AMH synthesis in our experimental animals that received stem cell injections, observing a concurrent increase in AMH synthesis. The observation of restored contralateral ovarian function along with elevated AMH synthesis by the differentiated stem cells collectively supports the idea that AMH may be important in mediating the observed restoration of ovarian function in these studies *(43–45, 47, 48)*. Together, these data suggest the feasibility of patients providing their own cells to restore ovarian function, synthesize bioidentical hormones, and restore fertility by neogametogenesis.

In such a clinical paradigm, autologous iPSCs from a patient may be differentiated into steroidogenic tissue to generate bioidentical reproductive hormones (progesterone, estradiol, testosterone, and AMH; Fig. 7). These patient-specific hormones could be used for hormone replacement therapy (HRT), while ovarian tissue and primordial gametes derived from these iPSCs may prove useful for *in vitro* egg maturation (IVM) in patients requiring ART. Such an IVM module could be compartmentalized in a patient-specific microfluidic chip for clinical applications (Fig. 7). We believe that these findings collectively afford us the opportunity to develop novel therapeutic options for cell-based therapies, especially relevant for women with premature ovarian failure (POF).

**Figure 7.**
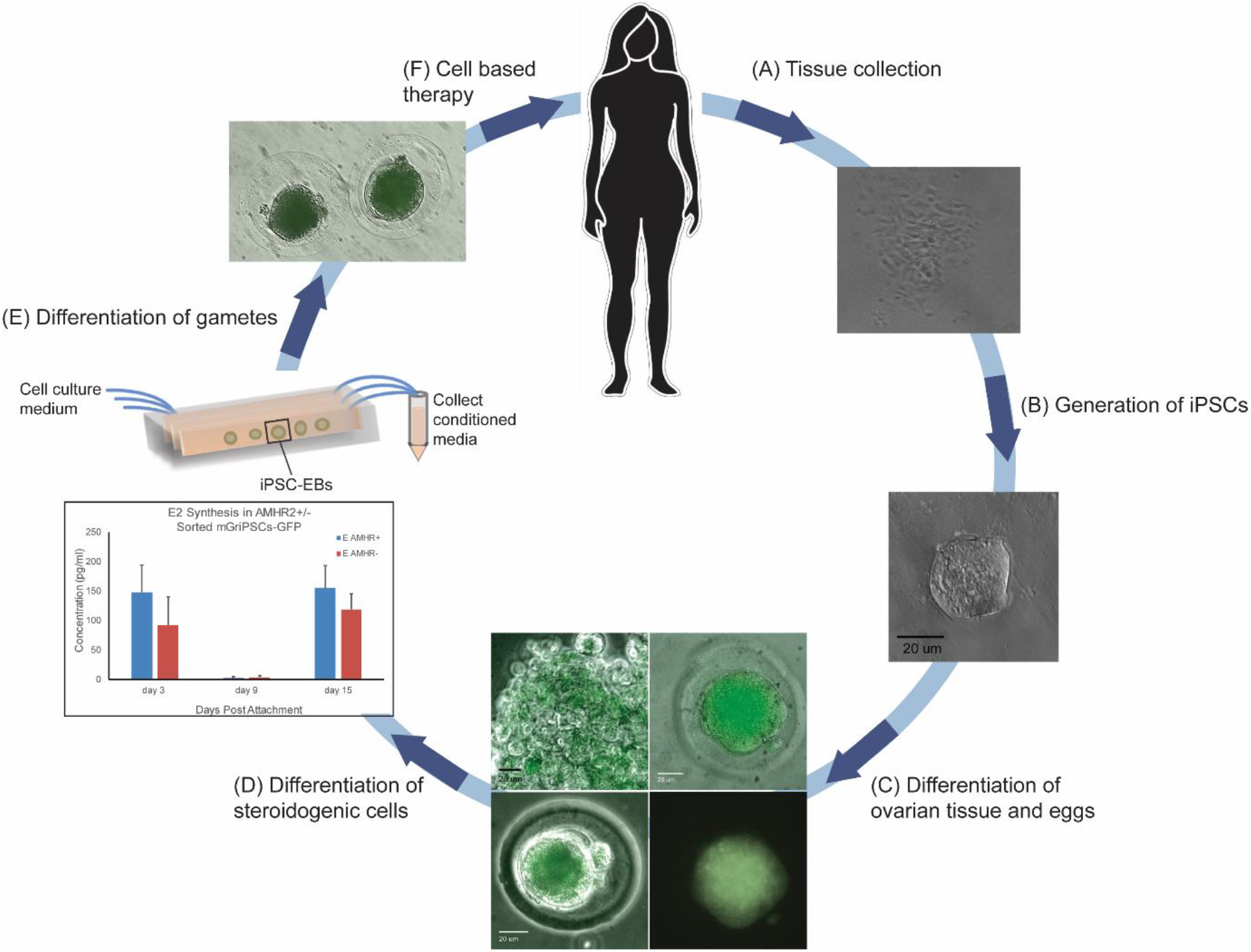
An autologous stem cell-based model for fertility and endocrine function restoration. A model for translational applications of this research. Somatic cells are collected from a patient (**A**) and converted into autologous iPSCs (**B**). These iPSCs are differentiated into ovarian steroidogenic and reproductive cells (**C**, **D**). Microfluidic chip technology may be utilized to create a sterile, continuously flowing environment, creating the ability to purify autologous bioidentical reproductive hormones from the conditioned media (**D**). The autologous oocytes (**E**) and endogenous steroid hormones (**D**) may then be collected and used to treat the same patients, ultimately restoring fertility and endocrine function (**F**).

We do acknowledge several limitations to our study. First is the use of granulosa cell-derived iPSCs. To determine whether our endocrine differentiation observations may be extended to other iPSC types, we have previously investigated fibroblast and amniocyte derived iPSCs as well as embryonic stem cells with similar results, although with a lower efficiency *(18, 20)*. We believe the higher efficiency of mGriPSCs in generation of ovarian endocrine tissue relates to the epigenetic memory of these cells. However, the observed ability to differentiate other iPSC types along this pathway potentially benefits a larger number of patients who may not have had the opportunity to attempt IVF and harvest granulosa cells prior to their chemotherapy or who have had oophorectomies and no longer have ovarian tissue present. Second, while we have shown that AMHR2+ mGriPSCs can rescue ovaries exposed to chemotherapy, we have not tested whether AMHR2+ mGriPSCs can be derived from ovaries that have already received a gonadotoxic insult. We have also not tested the ability to impact other gonadotoxic insults, such as radiation therapy. Finally, we have not thoroughly evaluated the normalcy of these stem cell-derived oocytes and pups. We hope to address these aspects in future work.

In sum, the ability to generate patient-specific bioidentical hormones and autologous oocytes would revolutionize therapeutic options for women with chemotherapy-related or idiopathic premature ovarian failure, who comprise up to 5% of the general population of women *(54)*. In the future, the generation of functional human gametes derived from iPSCs could prove a major advancement in translational fertility research.

## Materials and Methods

### Study design

The objective of this study was to explore the *in vivo* reproductive and steroidogenic potential of a novel line of mouse iPSCs (i.e. mGriPSCs) that were derived from somatic ovarian granulosa cells *(18)*. mGriPSCs were genetically labeled with GFP for lineage tracing and the resulting mGriPSCs-GFP were differentiated in media containing HFF. FACS was employed to purify a subpopulation of differentiated mGriPSCs-GFP that expressed AMHR2, an established ovarian marker. These AMHR2+ mGriPSCs-GFP were then orthotopically injected into the ovaries of subfertile nude mice, which had previously been exposed to gonadotoxic chemotherapy. In addition to assessing reproductive hormone synthesis and RNA and protein expression in ovarian tissues extracted from these mice, oocytes were also retrieved, assessed for GFP expression, and *in vitro* fertilized or activated with a calcium ionophore. In a second round of experiments, subfertile nude mice were again orthotopically injected with AMHR2+ cells and subsequently bred with wild type black 6 males. The resulting embryos and pups were assessed for morphology and GFP expression.

All supplies were purchased from Sigma Aldrich (St. Louis, Missouri), unless otherwise stated. All antibodies were purchased from Abcam (Cambridge, MA; Table S1) and all PCR primers were purchased from Thermo Fisher (Waltham, MA; Table S2). All protocols involving animals or using animal tissue have been approved by Brigham and Women’s Hospital Institution of Animal Care and Use Committee (IUCAC), detailed in protocol #2016N000367 or Dana Farber Cancer Institute IUCAC, detailed in protocol #15-047. All experiments were performed in accordance with relevant guidelines and regulations.

### Generation of mGriPSCs-GFP

Mouse granulosa cells were reprogrammed as previously described to generate mGriPSCs *(25)*. Infecting mGriPSCs with GFP allowed the mGriPSCs and resulting differentiated cells to be labeled and tracked throughout our experimental process. The GFP gene was transfected into 293T cells by combining the GFP construct, VSV-G, and delta 8.2 lentiviral packaging system with FuGENE and culture media. Using a fluorescent microscope (Nikon), the GFP signal was observed in transfected 293T cells. The viral containing culture media was harvested and mixed with 8 μg/ml polybrene to be fed onto healthy stem cell colonies. Resulting mGriPSCs-GFP were enzymatically dissociated with 0.05% trypsin-EDTA and GFP expression was confirmed by FACS and live fluorescent microscopy. Healthy mGriPSCs-GFP stem cell colonies were observed and stem cell status was verified by RT-PCR (Fig. S1) and immunocytochemistry (ICC), as well as an alkaline-phosphatase reaction kit to ensure pluripotency. Commercial stem cell antibodies, OCT4 (Abcam), SSEA-1 (Millipore), and NANOG (Abcam), were used for ICC verification. The RT-PCR was performed using the primers of stem cell markers *Oct4, Nanog, Gdf3*, and *Dnmt3b* to verify pluripotency of the mGriPSCs-GFP. Embryoid bodies (EBs) were then cultured as previously described *(18(18)*, with HFF added to EB culture media at various concentrations ranging from 1% to 5% to assess which concentration most effectively promoted differentiation of mGriPSCs to ovarian cell types. 1% HFF EB media was used to culture all mGriPSCs-GFP that would later be FACS-purified and injected into mice.

### Human Follicular Fluid (HFF) acquisition

The following protocol involving human participants was approved by Partners Human Research Committee (PHRC), the Institutional Review Board (IRB) of Partners HealthCare (protocol #2011P000795). All experiments were performed in accordance with relevant guidelines and regulations. Written informed consent was obtained from all participants. HFF was obtained from our institution’s IVF laboratory as discarded tissue from consenting patients. Oocyte retrievals were performed as part of the patients’ routine care. After transvaginal ultrasound-guided aspiration of oocytes as well as the follicular fluid was completed, discarded HFF was collected, anonymized, and pooled from different patients into one collection tube. Freshly obtained HFF was centrifuged at 1500 RPM for 5 min and supernatant was collected and passed through a 0.22 um filter to remove impurities and contaminants. A standard endocrine assay was used to determine concentration of E2, FSH, and LH in HFF samples. Filtered HFF samples were frozen at −80°C until necessary cell culture use.

### Fluorescence-activated cell sorting (FACS)

To further purify the subpopulation of presumptive ovarian and oocyte cells from the differentiated mGriPSCs-GFP cells, live FACS was employed, using the ovarian cell surface marker, AMHR2. To prepare for FACS, EBs were dissociated into single cells by exposing them to pre-warmed 0.05% trypsin-EDTA and passing them through a 40 um strainer. Single cells were subsequently stained with a mouse AMHR2 primary antibody (Abcam) for one hour, rinsed, and treated with anti-mouse Alexa Flour 488 secondary antibody (Life Technologies/Invitrogen, Carlsbad, CA). Using BD FACSAria multicolor high-speed sorter and FACSDiva version 6.1.2 software (BD Biosciences, Franklin Lakes, New Jersey), cells were separated into either AMHR2+ and AMHR2-groups. AMHR2+ sorted cells were either injected on the same day into experimental mice or plated on gelatin-coated plates and cultured in EB media for one week, while AMHR2-cells were only plated and cultured.

### Mouse gonadotoxic chemotherapy and intramuscular injection

Female B6.Cg-Foxn1^nu^/J homozygous nude mice were received from Jackson Laboratory (Bar Harbor, ME, Cat# 000819). Mice were randomized into four groups: 1) control, 2) chemotherapy with no stem cell injection, 3) chemotherapy with unsorted mGriPSCs injection, and 4) chemotherapy with sorted mGriPSCs injections. The control group contained four mice, while all other groups had three. Twelve days after arrival, six-week old mice were hyperstimulated with 5 IU of PMSG followed 48-hours later by 5 IU of hCG to synchronize the mice’s menstrual cycles. Three weeks later, premature ovarian insufficiency was induced by single intraperitoneal injections of busulfan (12 mg/kg) and cyclophosphamide (120 mg/kg). Control mice received 100 ul of vehicle (10% DMSO in PBS). After an additional three weeks, experimental mice received intramuscular injections of 2 million unsorted mGriPSCs, sorted mGriPSCs, or vehicle into the left thigh. Hormone synthesis was analyzed and compared between mice that received chemotherapy and controls. Teratomas were measured in mice that received unsorted injections, while no teratomas were detected in mice that received sorted injections.

### Mouse intraovarian cell injection

Mice were randomized into the same four groups as described above, but this time, each group comprised of five mice. Chemotherapy was administered in eight-week old nude mice as described above. One week after chemotherapy administration, mice underwent laparotomy with intraovarian injection of either sorted or unsorted mGriPSCs-GFP. The laparotomy was performed by placing the mice under isoflurane anesthesia, then infiltrating the ventral midline with a 1:1 mixture of lidocaine and bupivacaine. A midline anterior vertical laparotomy was made under sterile technique. In each case, the left ovary was gently elevated and injected with 2 million cells suspended in PBS using a 27-gauge needle. The ovary was then returned to the abdomen and the abdominal wall closed in two layers using suture. Mice received 72 hours of meloxicam for post-operative analgesia. Experiments were repeated twice.

### Mouse intraovarian cell injection, oocyte retrieval, and breeding

In a separate round of experiments, differentiated AMHR2+ mGriPSCs-GFP were orthotopically injected into both ovaries, as described above, of five eight-week old, immunocompromised homozygous nude mice (Jackson Laboratory, Cat# 002019), which are known to be subfertile. Five weeks after stem cell injections, two of these mice were hyperstimulated with PMSG and hCG, as described above. These mice were then sacrificed for oocyte retrieval and IVF, as described below. Eight weeks after stem cell injections, the remaining three stem cell-injected nude mice were mated with eight-week old, wild type Black 6 male mice (Jackson Laboratory, Cat# 000664). All three mice became pregnant, and two of the three pregnant mice were euthanized around embryonic day 18 in order to assess embryos for morphology and expression of GFP via confocal microscopy, IHC, and RT-PCR. Tail snips were performed on E18 embryos with samples snap frozen and stored in −80°C for subsequent nucleotide extraction and RT-PCR. The embryos were then fixed and immunostained for IHC, as described below in the “ICC/IHC” section. The third mouse was mated three times within the span of eight weeks and was followed to delivery each time. Tail snips were collected from live pups in order to assess GFP expression via IHC and RT-PCR.

### Oocyte retrieval, activation and *in vitro* fertilization

Oocytes were retrieved as previously described *(56)* and cultured in KSOM at 37°C and 5% CO_2_. Half of the retrieved oocytes were treated with hyaluronidase to strip the cumulus granulosa cells, followed by exposure to calcium ionophore A23187 in KSOM media overnight at 37°C in 5% CO_2_. The rest of the oocytes were treated with sperm extracted from the cauda epididymis of C57BL6/J mice. Using a phase-contrast light microscope (Zeiss, Oberkochen, Germany), oocytes were observed for 3 days for any signs of activation or fertilization. Oocytes, granulosa cells, and half of each ovary were snap frozen in liquid nitrogen for subsequent RNA extraction. The other half of the ovary was submerged in optimal cutting temperature (OCT) compound (Thermo Fisher), placed on dry ice to freeze, and stored in −80°C freezer for future IHC analysis.

### Processing of control ovarian tissues

Untreated C57BL6/J mice were sacrificed and their ovaries excised. Ovaries were fixed in cold 4% paraformaldehyde (PFA)/4% sucrose for 30 minutes, followed by immersion in 50%, 80%, 95%, and 100% ethanol for two hours each, 100% ethanol overnight, xylene for four hours, xylene overnight, 56°C paraffin for two hours twice, and finally embedded in paraffin. Serial sections of the ovary were cut in a cryostat (Thermo Fisher) at a thickness of 5 um and stained by hematoxylin and eosin (H&E) as well as immunostained for oocyte and ovarian markers listed below in the “ICC/IHC” section. The serial sectioning of control ovaries was necessary to provide comparative staining for our experimental cells and tissues, displaying ovarian and oocyte markers.

### Immunocytochemistry/Immunohistochemistry (ICC/IHC)

Pre-warmed 0.05% trypsin-EDTA was used to dissociate EBs so that they could be subsequently reattached to gelatin-coated slides as a more ideal visual monolayer of cells. Larger tissues, such as ovaries, or E18 embryos were first fixed in 4% PFA/4% sucrose (30 minutes for smaller tissues such as ovaries, overnight for whole embryos), followed by immersion in 10%, 20%, and 30% sucrose in PBS, with tissues or embryos immersed until they sank to the bottom of the vial. Tissues and whole embryos were then mounted in OCT, frozen on dry ice, and cut in a cryostat with serial sections of 5 um thickness. Once placed on slides, cells or prepared histological tissue or embryo sections were fixed in 4% PFA/4% sucrose, immunostained, and visualized as described previously^18^. Briefly, cells and tissues were blocked with 2% donkey serum, 10 mg/mL bovine serum albumin, and 1% Triton-X for 30 minutes. Primary antibodies were then applied at a concentration of 1:500 for two hours at room temperature. Secondary antibodies were applied at a concentration of 1:1000 for one hour at room temperature, followed by application of 4’,6-diamidino-2-phenylindole (DAPI) for 10 minutes at room temperature to visualize the nuclei. The primary antibodies for ovarian markers, AMHR2, CYP19a1, FOXL2, FSHR, GJA1, INHB (Santa Cruz, Dallas, TX), and primary antibodies for oocyte markers, DAZL, DDX4, ZP1, and ZP2 (Santa Cruz), were used for this analysis. For the cell count (Fig. 1 VI), positively stained cells were counted in 7 distinct fields of view per slide, as described previously *(18)*.

### RT-PCR and Gel Electrophoresis

After RNA was extracted using commercially available kits (Qiagen, Germantown, MD), cDNA was synthesized via a qScript cDNA Synthesis kit (Quanta Biosciences, Gaithersburg, MD). The cDNA, DNA polymerase, and master mix (Promega, Madison, WI) were combined with corresponding primers for ovarian markers *Amhr*2, *Cyp19a1*, *Fshr*, *Gja1*, and *Amh*, and oocyte markers *Ddx4*, *Boule*, and *Blimp-1*, with *β-actin* as a positive control. Cycling conditions were 95°C for 3 minutes, 35 repetitions of (95 °C for 1 minute, 58.5 °C for 1 minute, and 72 for 1 minute) and 72 °C for 10 minutes in a thermocycler (Bio-Rad). Amplified products were separated on 1.0% agarose gel electrophoresis (Thermo Fisher) to qualitatively analyze the expression of tested biomarkers.

### RT-qPCR

Homogenized and lysed tissue and cell samples were processed for RNA extraction as described above. After cDNA synthesis, 2 ng of cDNA was used in each qPCR reaction well. Primers for *Zp1, Boule, Ddx4, Gja1, Inhb, Cyp19a1*, and *Amh* were used for qPCR of ovary tissues. Power SYBR Green reaction mix (Applied Biosystems, Foster City, CA) was used for qPCR reactions. A QuantStudio 3 (Applied Biosystems) real-time thermocycler was used with the cycling conditions of 50 °C for 2 minutes, 95 °C for 10 minutes, and 60 repetitions of 95 °C for 1 minute and 60 °C for 2.5 minutes. Fold changes in gene expression were measured using the ∆∆Ct technique with *β-actin* as the endogenous control gene for relative quantification. qPCRs for the ovarian tissue used ovaries that were not treated with chemotherapy nor injected with stem cells as the referent control against which fold change in gene expression was compared in the treated ovaries.

### ELISA and hormone analysis

Estradiol, progesterone, and AMH concentrations were analyzed using commercially available ELISA kits (Abnova, Zhongli, Taiwan). Three media samples from each cell culture condition were collected. Additionally, serum from the four different mouse treatment groups were collected and analyzed at three timepoints (baseline, post-chemotherapy, and pre-sacrifice).

### Statistical analysis

All data was compared and analyzed using Mann-Whitney U tests, with p<0.05 for significance.

## Acknowledgements

The authors would like to thank Dr. Robert Barbieri and Dr. Mark Hornstein for editorial comments, Maya Seshan for editorial assistance, and Mazhar Chaudhry for assistance with endocrine analyses.

## Author contributions

K.E. and R.M.A. developed the research design. E.R.D., E.S.G., K.E., G.C., and R.M.A. wrote the manuscript. A.G., A.M., E.R.D., K.E., K.U.D., N.W.N., N.D., and R.M.A. performed the experiments and collected data. A.M., K.U.D., N.W.N., and R.M.A. analyzed the data.

## Data and materials availability

All data associated with this study are available in the main text or the supplementary materials.

