## Supplementary Figure 1 for "iPSC-Derived Ovarian Tissue Restores Ovarian Function in Subfertile Mice and After Gonadotoxic Chemotherapy"

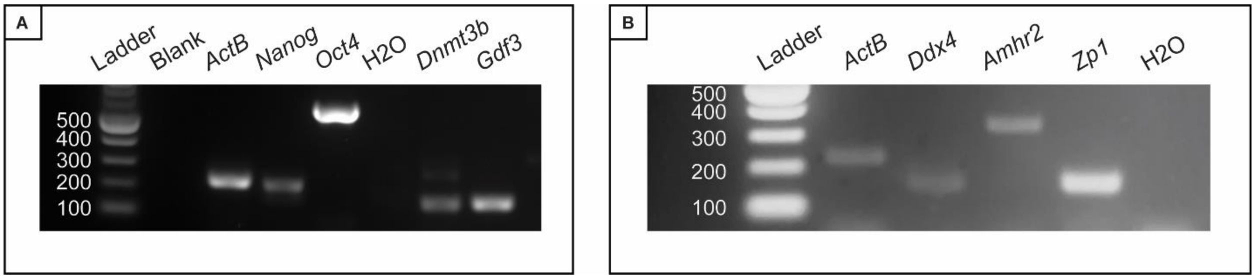


**Fig. S1. mGriPSCs-GFP express stem cell markers and mouse oocytes express *Amhr*.** RT-PCR analysis shows that stem cell colonies of mGriPSCs-GFP maintain expression of pluripotency markers *Nanog*, *Oct4*, *Dnmt3b*, and *Gdf3*. RT-PCR also shows that oocytes collected from control mice express ovarian marker *Amhr* and oocyte markers *Zp1* and *Ddx4*.
