## Supplementary Figure 2 for "iPSC-Derived Ovarian Tissue Restores Ovarian Function in Subfertile Mice and After Gonadotoxic Chemotherapy"

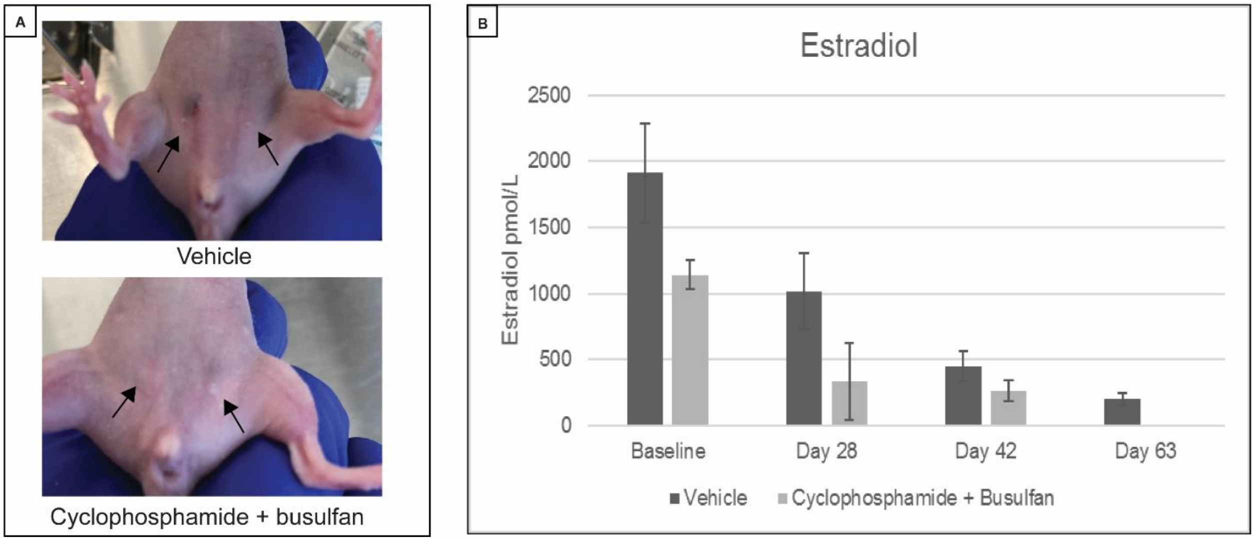


**Fig. S2. Chemotherapeutic agents decrease estradiol production, while injection of unsorted stem cells results in teratoma formation.** Busulfan and Cyclophosphamide injections result in breast atrophy in nude mice compared to control mice receiving vehicle injections (A). Estradiol synthesis decreases in mice receiving Busulfan and Cyclophosphamide compared to control mice, as measured by ELISA over 4 timepoints (B).
