## Supplementary Figure 3 for "iPSC-Derived Ovarian Tissue Restores Ovarian Function in Subfertile Mice and After Gonadotoxic Chemotherapy"

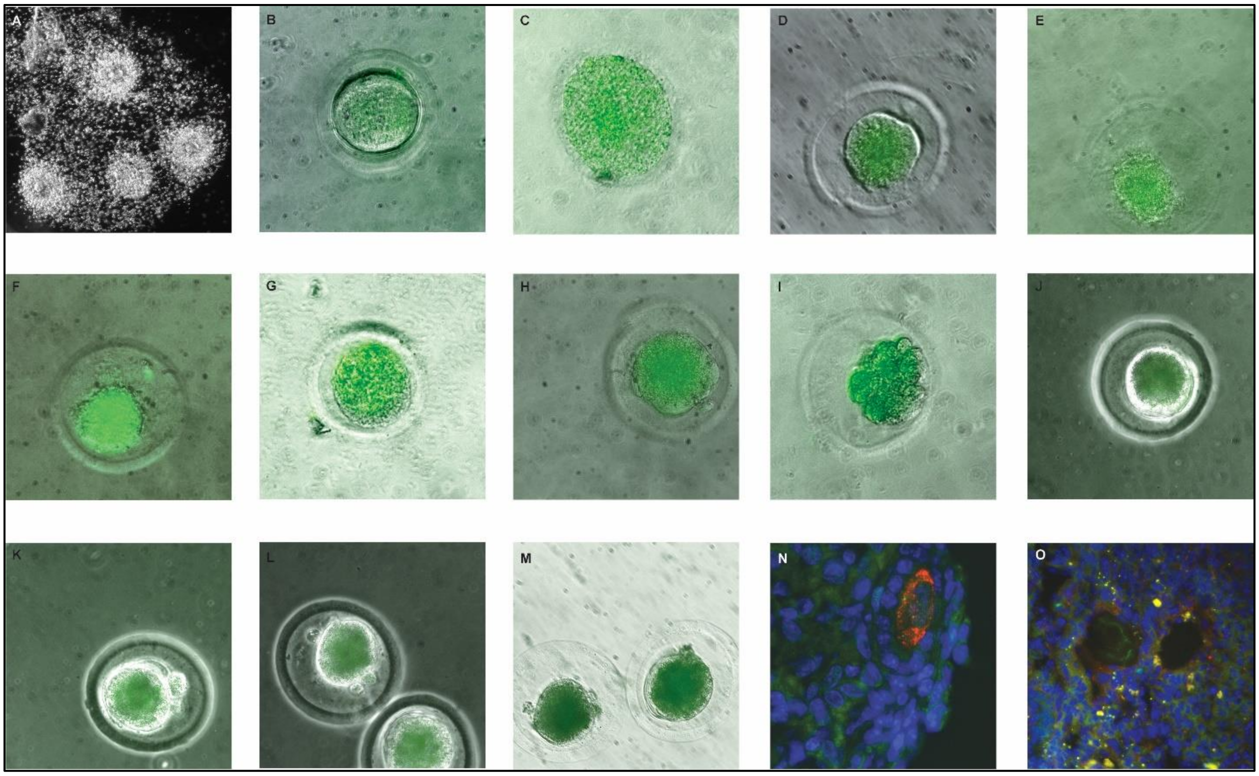

**Fig. S3.** ***De novo*** **generation of mature stem cell-derived oocytes.** Phase contrast image of oocytes and surrounding granulosa cells retrieved from nude mice injected with sorted mGriPSCs-GFP (A). Retrieved oocytes express GFP (B-M) and are fertilized with mouse sperm (D-I) or activated by a calcium ionophore (A23817) (J-M). IHC on sorted mGriPSCs-GFP injected ovaries shows co-localization of DAZL (N) and AMHR (M) with GFP in the ovarian follicles of treated mice.
