## Supplementary Figure 4 for "iPSC-Derived Ovarian Tissue Restores Ovarian Function in Subfertile Mice and After Gonadotoxic Chemotherapy"

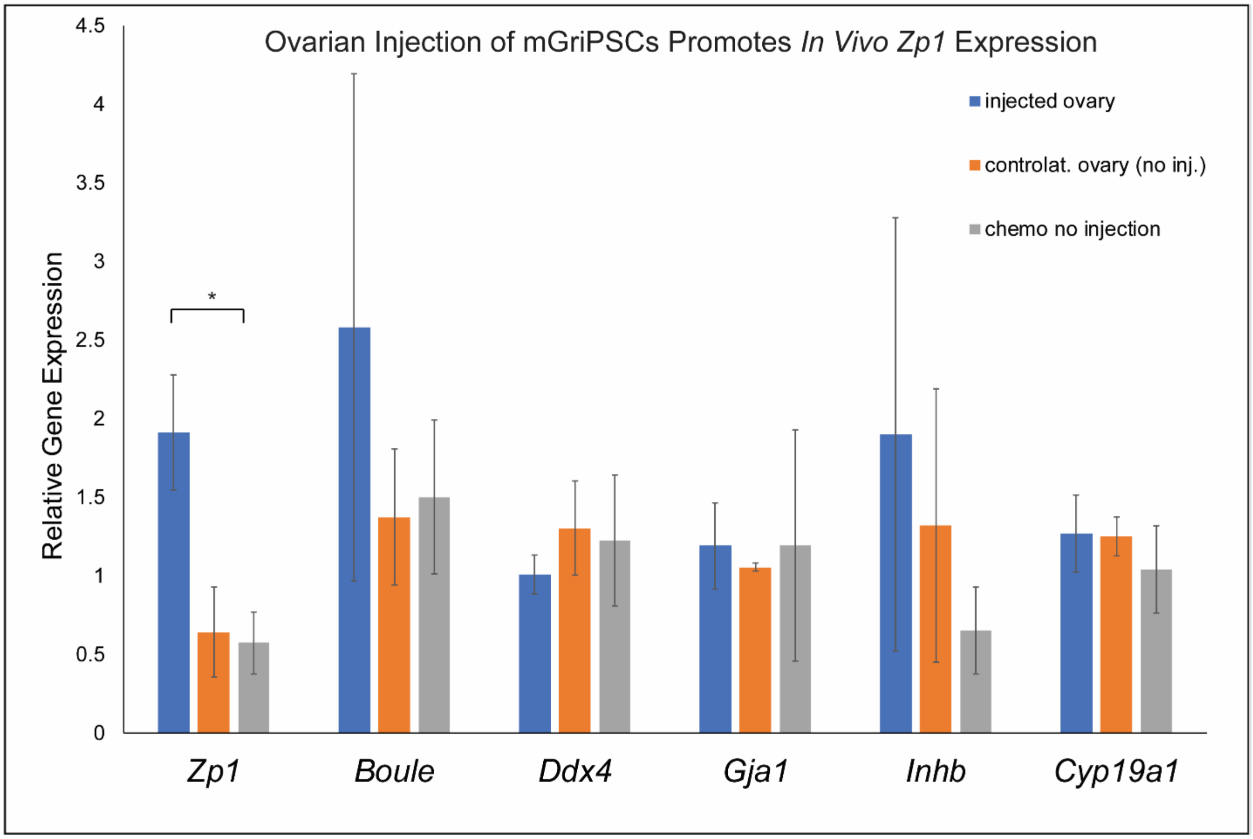


**Fig. S4. Ovarian and oocyte gene expression in treated mouse ovaries.** RT-qPCR analysis shows that ovaries exposed to chemotherapy and injected with sorted mGriPSCs-GFP expressed higher levels of *Zp1* compared to ovaries contralateral to the injected ovary and ovaries from mice that received chemotherapy but no stem cell injections (n=3, Mann-Whitney U test, $p<0.05$, data represented as mean ± SEM gene expression fold change as compared with control ovary gene expression).
