## Supplementary Figure 5 for "iPSC-Derived Ovarian Tissue Restores Ovarian Function in Subfertile Mice and After Gonadotoxic Chemotherapy"

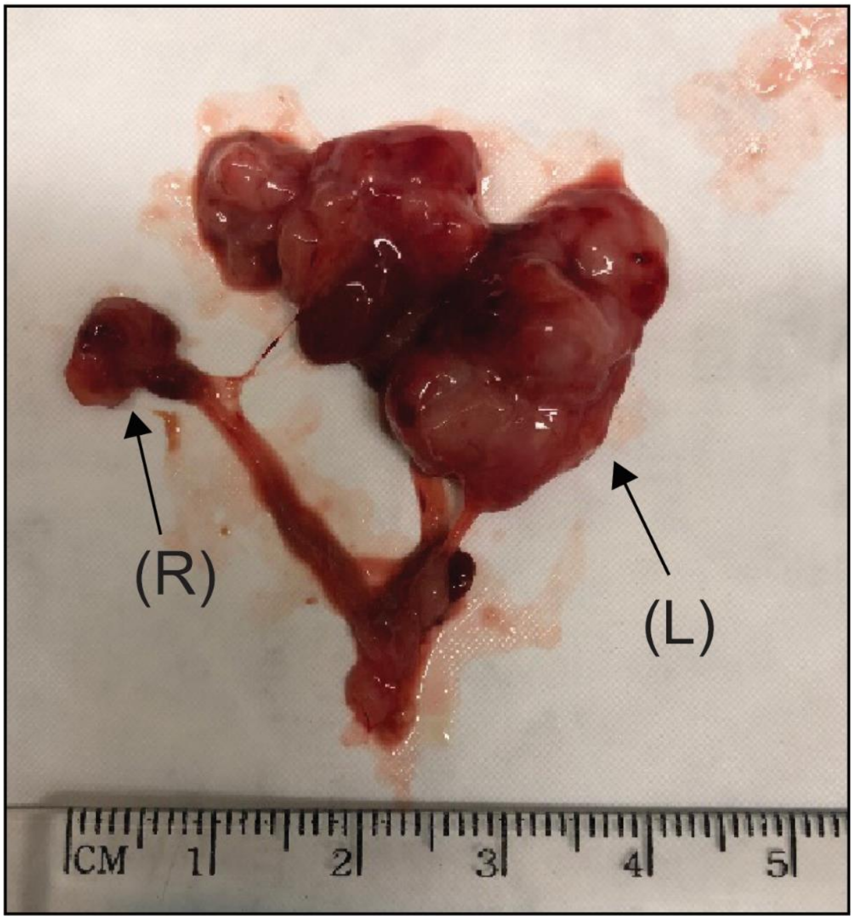


**Fig. S5. Teratoma formation from injections of unsorted mGriPSCs into the left ovary.** The teratoma formation inhibited oocyte retrieval.
