## Supplementary Figure 6 for "iPSC-Derived Ovarian Tissue Restores Ovarian Function in Subfertile Mice and After Gonadotoxic Chemotherapy"

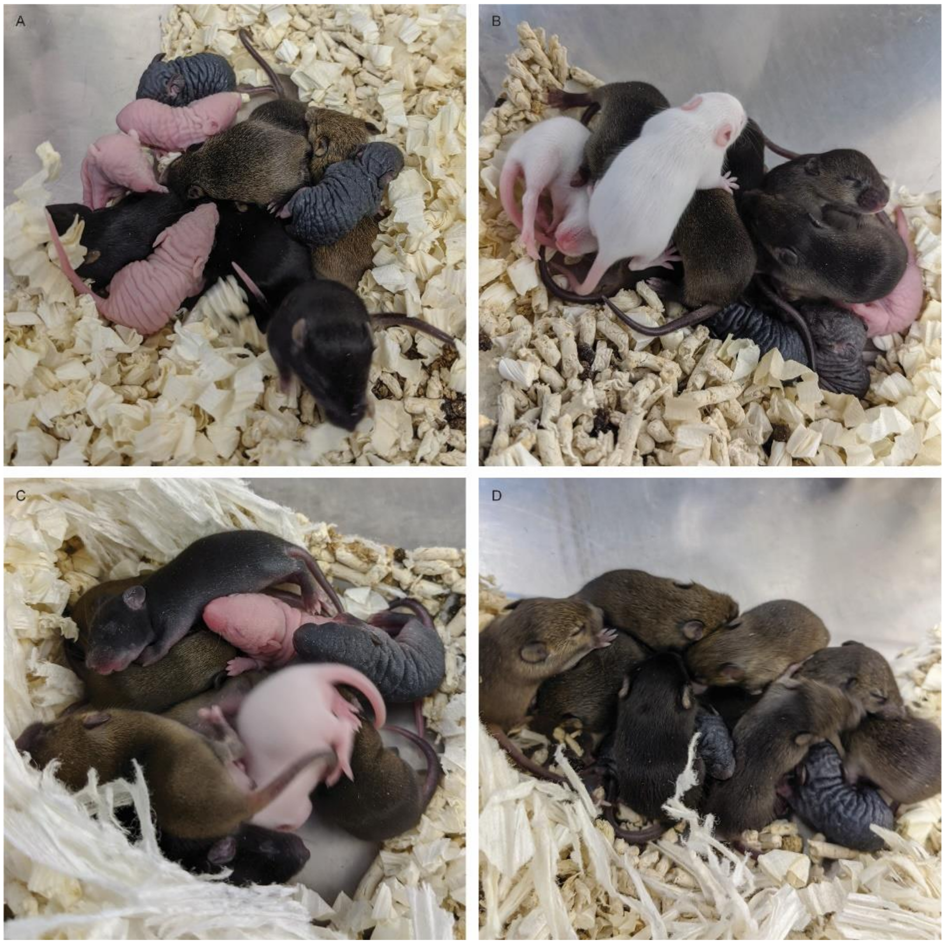


**Fig. S6. Four litters of F2 pups after the mating of F1 pups.** Four female F1 pups were mated with two F1 male pups. All four females gave birth to litter sizes of 11 (A), 11 (B), 10 (C), 9 (D) pups. The F2 litters (A-D) varied in skin color, coat, and coat color.
