## Supplementary figures and images for "iPSC-Derived Ovarian Tissue Restores Ovarian Function in Subfertile Mice and After Gonadotoxic Chemotherapy"

### Supplementary Figure 7

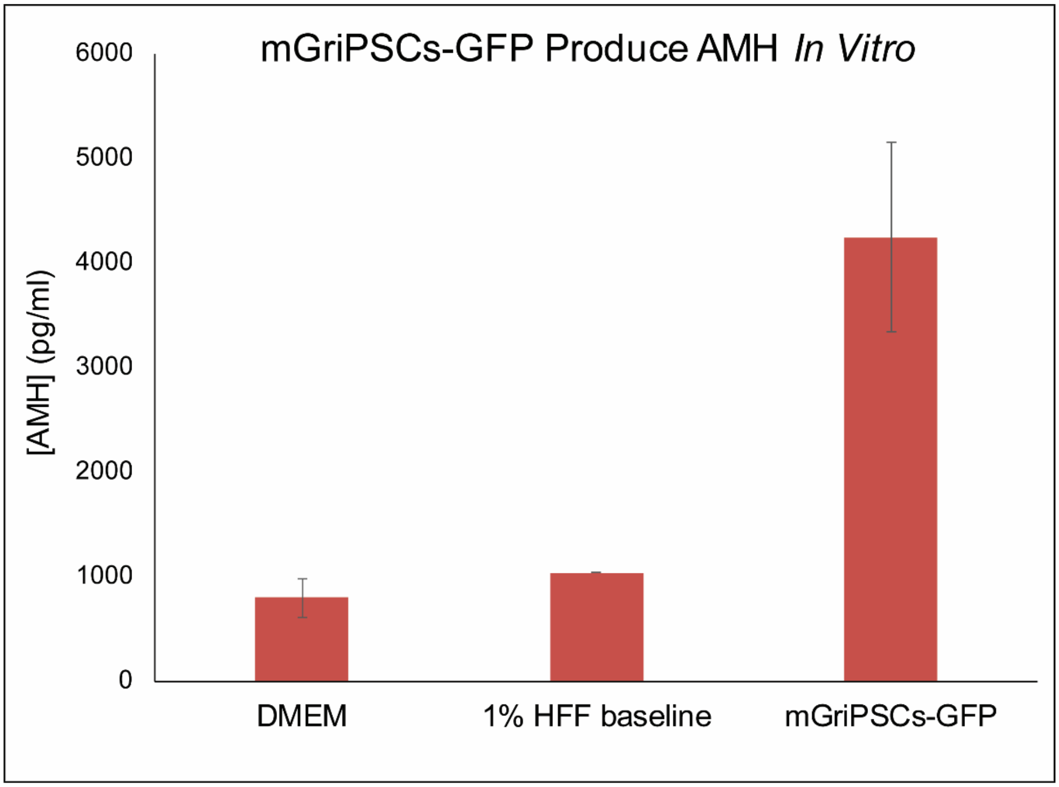


**Fig. S7.** Attached differentiated mGriPSCs synthesized AMH *in vitro*.
