## Supplementary Table 1 for "iPSC-Derived Ovarian Tissue Restores Ovarian Function in Subfertile Mice and After Gonadotoxic Chemotherapy"

**Table S1.** Antibodies used for immunolabeling.

| **Antibody** | **Company** | **Cat. #** |
|---|---|---|
| AMH | Santa Cruz | Sc-6886 |
| AMHR2 | Abcam | Ab64762 |
| BOULE | Abcam | Ab104491 |
| CYP19a1 | Abcam | Ab35604 |
| DAZL | Abcam | Ab34139 |
| FSHR | Santa Cruz | Sc-7798 |
| FOXL2 | Abcam | Ab5096 |
| GJA1 | Abcam | Ab11370 |
| INH β-A | Santa Cruz | Sc-166503 |
| MVH/DDX4 | Abcam | Ab13840 |
| NANOG | Abcam | Ab106465 |
| OCT4 | Abcam | Ab18976 |
| SSEA1 | Millipore | MAB4301 |
| ZP1 | Santa Cruz | Sc-23706 |
| ZP2 | Santa Cruz | Sc-32752 |
