## Supplementary Table 2 for "iPSC-Derived Ovarian Tissue Restores Ovarian Function in Subfertile Mice and After Gonadotoxic Chemotherapy"

**Table S2.** PCR primer sequences.

| Gene | Forward Sequence | Reverse Sequence |
|---|---|---|
| *Actb* | TGTTACCAACTGGGACGACA | CCATCACAATGCCTGTGGTA |
| *Amh* | GCAGTTGCTAGTCCTACATC | TCATCCGCGTGAAACAGCG |
| *Amhr2* | GTATCCGCTGCCTCTACAGC | AGCCTGGCTCATCACTGTCT |
| *Blimp1* | CTTCTCTTGGAAAAACGTGTGG G | TCATATCAGCGTCCTCCATG |
| *Boule* | TTGTCAATCCGGCCATTTGC | CCGTATCTTGGGCCACTTGT |
| *Cyp19a 1* | CAAAGCACGCTGCAAATACCA | GGCCAAATGTGTCTTCCAGT |
| *Dnmt3B* | GCGCAGCGATCGGCGCCGGAGA T | CATACCCGGTGGCACCCTGTTCTTC AGTCA |
| *Foxl2* | GCTATTTAGGTGACACTATAGTC ATAGCCAAGTTCCCGTTC | TTGTAATACGACTCACTATAGGGCC AGGAGTTGTTGAGGAA |
| *Fshr* | AGTTGCATGGCATGTGTGAT | CATCACTGGGAACACCACG |
| *Gdf3* | ACCTTTCCAAGATGGCTCCT | CCTGAACCACAGACAGAGCA |
| *Gfp* | CTCGCTTGTCGGCCATGATA | CACGACTTCTTCAAGTCCGC |
| *Gja1* | AACCTTGACTTCCGAGCAGA | CTTGGGGAACAAAGGAATCA |
| *Inh β-A* | GCTATTTAGGTGACACTATAGA TCATCACCTTTGCCGAGTC | TTGTAATACGACTCACTATAGGGCT TCTTCCCATCTCCATCCA |
| *Mvh/Dd x4* | GAGAACACATCTACAACTGGTG G | CCTCGCTTGGAAAACCCTCT |
| *Nanog* | CAGGTGTTTGAGGGTAGCTC | CGGTTCATCATGGTACAGTC |
| *Oct4* | CACGAGTGGAAAGCAACTCA | CTGGGAAAGGTGTCCCTGTA |
| *Sry* | GCTATTTAGGTGACACTATAGA TGGAGGGCCATGTCAAG | TTGTAATACGACTCACTATAGGGAA CAGGCTGCCAATAAAAGC |
| *Zp1* | CCCTGAGATTGGGTCAGCG | AGAGCAGTTATTCACCTCAAACC |
