## Supplementary Table 3 for "iPSC-Derived Ovarian Tissue Restores Ovarian Function in Subfertile Mice and After Gonadotoxic Chemotherapy"

**Table S3.** List of abbreviations

| **Name** | **Abbreviation** |
|---|---|
| 4’,6-diamidino-2-phenylindole | DAPI |
| anti-Müllerian hormone | AMH |
| anti-Müllerian hormone receptor 2 | AMHR2 |
| Assisted reproductive technology | ART |
| Aromatase | CYP19A1 |
| DEAD-box helicase 4 | DDX4 |
| Deleted in zoospermia-like | DAZL |
| Embryonic stem cells | ESCs |
| Enzyme-linked immunosorbent assay | ELISA |
| Fluorescent activated cell sorting | FACS |
| Follicle-stimulating hormone | FSHR |
| Forkhead box protein L2 | FOXL2 |
| Gap junction alpha-1 | GJA1 |
| Gonadotropin releasing hormone | GnRH |
| Granulosa cell | GC |
| Green fluorescent protein | GFP |
| Heat-inactivated fetal bovine serum | HI FBS |
| Hematoxylin and eosin | H&E |
| Hormone replacement therapy | HRT |
| Human chorionic gonadotropin | hCG |
| Human follicular fluid | HFF |
| Immunocytochemistry | ICC |
| In vitro fertilization | IVF |
| Induced pluripotent stem cells | iPSCs |
| Inhibin β-A | INHBA |
| Institution of Animal Care and Use Committee | IUCAC |
| Knock-out serum replacement | KSOR |
| Mouse embryonic fibroblasts | MEFs |
| Mouse granulosa cell derived-induced pluripotent stem cells | mGriPSCs |
| Non-essential amino acids | NEAA |
| Optimizing cutting temperature | OCT |
| Phosphate-buffered saline | PBS |
| Polymerase chain reaction | PCR |
| Potassium simplex optimized medium | KSOM |
| PR domain zinc finger protein-1 | BLIMP-1 |
| Pregnant mare serum gonadotropin | PMSG |
| Premature ovarian failure | POF |
| Premature ovarian insufficiency | POI |
| Stage-specific embryonic antigen-1 | SSEA-1 |
| Virus-G glycoprotein | VSV-G |
| Zona pellucida | ZP |
